# Pangenome-wide and molecular evolution analyses of the *Pseudomonas aeruginosa species*

**DOI:** 10.1101/020305

**Authors:** Jeanneth Mosquera-Rendón, Ana M. Rada-Bravo, Sonia Cárdenas-Brito, Mauricio Corredor, Eliana Restrepo-Pineda, Alfonso Benítez-Páez

## Abstract

**Background:** Drug treatments and vaccine designs against the opportunistic human pathogen *Pseudomonas aeruginosa* have multiple issues, all associated with the diverse genetic traits present in this pathogen, ranging from multi-drug resistant genes to the molecular machinery for the biosynthesis of biofilms. Several candidate vaccines against *P. aeruginosa* have been developed, which target the outer membrane proteins; however, major issues arise when attempting to establish complete protection against this pathogen due to its presumably genotypic variation at the strain level. To shed light on this concern, we proposed this study to assess the *P. aeruginosa* pangenome and its molecular evolution across multiple strains.

**Results:** The *P. aeruginosa* pangenome was estimated to contain more than 16,000 non-redundant genes, and approximately 15% of these constituted the core genome. Functional analyses of the accessory genome indicated a wide presence of genetic elements directly associated with pathogenicity. An in-depth molecular evolution analysis revealed the full landscape of selection forces acting on the *P. aeruginosa* pangenome, in which purifying selection drives evolution in the genome of this human pathogen. We also detected distinctive positive selection in a wide variety of outer membrane proteins, with the data supporting the concept of substantial genetic variation in proteins probably recognized as antigens. Approaching the evolutionary information of genes under extremely positive selection, we designed a new Multi-Locus Sequencing Typing assay for an informative, rapid, and cost-effective genotyping of *P. aeruginosa* clinical isolates.

**Conclusions:** We report the unprecedented pangenome characterization of *P. aeruginosa* on a large scale, which included almost 200 bacterial genomes from one single species and a molecular evolutionary analysis at the pangenome scale. Evolutionary information presented here provides a clear explanation of the issues associated with the use of protein conjugates from pili, flagella, or secretion systems as antigens for vaccine design, which exhibit high genetic variation in terms of non-synonymous substitutions in *P. aeruginosa* strains.

## Background

Humans are frequently infected by opportunistic pathogens that take advantage of their compromised immunological status to cause persistent and chronic infections. The Gram-negative bacterium *Pseudomonas aeruginosa* is one of those recurrent human pathogens. *P. aeruginosa* remains one of the most important pathogens in nosocomial infections, and it is often associated with skin, urinary tract, and respiratory tract infections [1]. Respiratory tract infections are of major relevance in cystic fibrosis patients, given that *P. aeruginosa* deeply affects their pulmonary function, causing life-threatening infections [2]. One of the better-known adaptive resistance mechanisms of *P. aeruginosa* to evade either the host immune response and drug therapy is its ability to form biofilms. The *Pseudomonas aeruginosa* biofilm is an extremely stable capsule-like structure constituted primarily of polysaccharides, proteins, and DNA, in which PsI exopolysaccharide seems to be a key player for biofilm matrix stability [3]. Quorum sensing signals promote the formation of *P. aeruginosa* biofilms, which minimizes the entry of antimicrobial compounds inside bacterial cells and hinders the recognition of pathogen-associated molecular patterns (PAMPs) by the host immune system [4]. Consequently, current treatments against *P. aeruginosa* fail to resolve infections before tissue deterioration occurs. To address this concern, more efficient alternatives to abolish *P. aeruginosa* infections have produced promising but not definitive results. Accordingly, several candidate *P. aeruginosa* vaccines have been developed by targeting outer membrane proteins (Opr), lipopolysaccharides (LPS), polysaccharides (PS), PS-protein conjugates, flagella, pili, and single or multivalent live-attenuated cells [5–9]. However, major issues in the development of a successful *P. aeruginosa* vaccine arise from the probable genotypic variation at the strain level, making *P. aeruginosa* a presumably antigenically variable organism. Results supporting this assumption have been reported, yielding genetic information from the *P. aeruginosa* genome. For example, genetic variability explored in multiple *P. aeruginosa* isolates from different regions of the world indicated that *pcrV*, a member of the type III secretion system, exhibits limited genetic variation in terms of non-synonymous substitutions [10]. Although this type of analysis is informative, it provides only a very limited view of the genetic and evolutionary processes occurring at the genome level in *P. aeruginosa* and does not completely explain the failure to design and develop a successful vaccine against this human pathogen. Although antigen selection to design a *P. aeruginosa* vaccine is not a reported problem [11], to date, no genomic studies have correlated antigen genetic structure and variation with the effectiveness of antibody immunotherapy or vaccines, the efficacy of which remains elusive [11]. Moreover, enormous variation in the response against *P. aeruginosa* immunogenic proteins in patients with *P. aeruginosa* infections [12] could indicate that genetic factors from the pathogen and/or host could be responsible for the incomplete efficacy of candidate vaccines tested. In this fashion, this study aimed to i) better understand the genome structure and genetic variation exhibited by *Pseudomonas aeruginosa¸* ii) link the genome variation information with past and future *P. aeruginosa* vaccine designs, and iii) present and validate new molecular markers for Multi-Locus Sequence Typing (MLST) based on the study of genes exhibiting a higher ratio of non-synonymous over synonymous substitution rate. To achieve these aims, a combined pangenome-wide and molecular evolution analysis was performed using up-to-date and genome-scale genetic information publicly available in the Pathosystems Resource Integration Center (PATRIC) database [13].

## Results and Discussion

### Defining the Pseudomonas aeruginosa pangenome

A total of 181 genomes of *P. aeruginosa* strains were obtained through the public PATRIC database (see methods and Additional File 1). The preliminary analysis of the *P. aeruginosa* genome size variability is shown in Table 1. The *P. aeruginosa* chromosome contains 6,175 genes on average, with a distribution ranging from 5,382 to 7,170 genes per genome, indicating a variation of 13-16% in terms of gene content among all strains analysed. By using the genome-centred approximation to define the *P. aeruginosa* pangenome (see methods), a total of 16,820 non-redundant genes were retrieved from those 181 genomes analysed. Almost one-third of the full set of genes constituting the *P. aeruginosa* pangenome, 5,209 genes (31%), were found to be uniquely present, meaning that every strain approximately contributes 29 new genes to the *Pseudomonas aeruginosa* pangenome on average. Initially, these data fit well with a theoretical number of strain-specific new genes added to the pangenome when a new strain genome was sequenced, 33 for the *Streptococcus agalactiae* pangenome [14]. However, for a more precise calculation of genomic and functional features of the *P. aeruginosa* pangenome, we performed general methods described by Tettelin and co-workers to define bacterial pangenomes [15]. After an iterative and combinatorial process, our observed data was plotted as rarefaction curves following Heaps’ law (Figure 1A). Further information was extracted from the pangenome analysis regarding gene categorization. The core genome or extended core of genes was characterized as the set of genes present in all or almost all genomes analysed; in this manner, we established that the *P. aeruginosa* core genome contains approximately 2,503 genes that are present in all 181 genomes studied, and they account for 15% of the pangenome. The graphical representation of the discovery rate for new genes at the core genome across the iterative analysis of *P. aeruginosa* strains for pangenome reconstruction is shown in Figure 1B. We analyzed such data with power law (n = κN^-α^) finding and averaged alpha parameter of 2.36 ± 0.49 (CI = 2.27 to 2.46) indicating the *P. aeruginosa* pangenome is closed according to proposed postulates of Tettelin and co-workers [15]. Moreover, interpretation of these data with the exponential regression allowed to estimate and horizontal asymptote (*θ*) 10 genes ± 2.99, indicating a small but finite number of new genes expected to be discovered with the study of new *P. aeruginosa* studies. A preliminary analysis regarding the distribution of some genes involved in lung infections like the biofilm-associated (*mifS*, *mifR*, *bamI*, *bdlA*, *bfiS*, and *bfmR*) and antibiotic resistance genes (*oprM, ampC, ampD*, and PIB-1), and functionally annotated in the Pseudomonas Genome Database [16], has revealed that these genetic entities are present between 95 to 100% of the strains studied here. This indicates that such functions, and pathogenicity by extension, are encoded into the core genome of *P. aeruginosa*.

**Figure 1.**
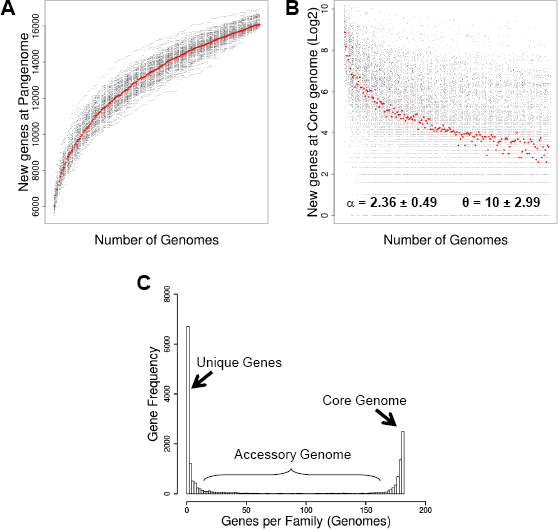
The *P. aeruginosa* pangenome. A – Rarefaction curve of the 200 different pangenomes calculated from random combinations of strains. Iterations and combinations are shown as the dots cloud indicating the total number of non-redundant genes included in the pangenome as genomes are included in the analysis. Red filled circles indicate the median of each iteration. B – Decay function for new genes discovered during pangenome reconstruction. Iterations and combinations are shown as dots cloud indicating the number of new genes incorporated to core genome. Red filled circles indicate the median of each iteration. The power law alpha parameter shown inside the plot is the average of such values retrieved individually in each iteration after fitting ± sd. The theta (*θ*) value was calculated from the horizontal asymptote where the exponential regression converges. C-Histogram for the prevalence of different gene families of the pangenome. The 16,820 non-redundant gene families determined to be present in the *P. aeruginosa* pangenome were distributed according to their frequency across all strains analysed. Three gene categories are clearly distinguished, highlighting the core genome (gene families present in all strains analysed), the unique genes (genes present in only one strain), and the accessory genome (gene families exhibiting a variable frequency).

**Table 1.** Main features of the *Pseudomonas aeruginosa* pangenome

| Features analysed | *P. aeruginosa* pangenome |
| --- | --- |
| Genomes | 181 |
| Total genes | 1,117,803 |
| Average genome size | 6,175 genes |
| Pangenome size (non-redundant genes) | 16,820 |
| Core genome | 2,503 genes |
| Accessory genome | 9,108 genes |
| Unique genes | 5,209 |
| Average unique genes/strain | 16 |
| Gene families under positive selection | 233 |

The set of genes, which were not included in the core genome or were unique (present in 1 genome), were referred to as the accessory genome; it included the 54% of genes found in the *P. aeruginosa* pangenome (Table 1). Interestingly, when we plotted the frequency of all pangenome genes present in different strains/genomes analysed (Figure 1C), we found a similar distribution to that reported by Lapierre and Gogarten when they estimated the pangenome for more than 500 different bacterial genomes [17]. This distribution plot clearly demonstrated the characteristic distribution and frequency of different groups of the above-stated genes. In general terms, the *P. aeruginosa* pangenome exhibits a high level of genome variability, whereby only 40% (2,503/6,175) of its genome is constant, on average. Thus, the remaining 60% of *P. aeruginosa* genome is presented as a variable piece of DNA composed of a wide repertoire of genes and molecular functions. A very recent study has partially characterized the *P. aeruginosa* pangenome using a total of 20 different human- and environmental-derived strains. Their numbers in terms of average genome size and ORFs per strains are very close to those we show in the present study. However, they estimate the *P. aeruginosa* pangenome to have 13,527 with more than 4,000 genes catalogued as the core genome [18]. The pangenome estimated in our study exceeds by more than 3,000 genes to that reported by Hilker and co-workers as well as to that reported by Valot and co-workers [19]. This is totally expected given that the more genomes analyzed, the more probability to discover new genes, an assumption that is clearly exemplified in the Figure 1A. Conversely, the core genome appear to be negatively affected by addition of new strains because the probability of sharing genes among strains decreases as new strains are incorporated to the study sample. This parameter intuitively is directly dependant of the number of strains used to calculate the core genome and their clonal relationship, which could strongly reduce gene diversity in the pangenome. Given that the multi-strain, iterative and combinatorial process used here to estimate the *P. aeruginosa* pangenome has produced a closed pangenome, we proposed that core genome for *P. aeruginosa* is composed of approximately 2,500 genes. This number is notably lower than those proposed in very recent studies aiming the characterization of the *P. aeruginosa* pangenome as well [18–20]. However, none of those studies have produced a proper metrics indicating that their proposed pangenomes are closed. Therefore, our data represent the most accurate characterization of the *P. aeruginosa* pangenome supported in the analysis of more than 180 different strains throughout iterative and combinatorial approaches. Moreover, the metrics presented in here is very close to that early characterized for *Escherichia coli*, for which a core genome was defined to account 2,200 genes [21].

Subsequently, we proceeded to perform a functional analysis with the full set of genes uniquely presented as well as other set of genes categorized by frequency in the *P. aeruginosa* pangenome. As a consequence, the nucleotide sequences of genes found to be present only in one *P. aeruginosa* strain were translated to amino acid sequences and then submitted to the Kyoto Encyclopedia of Genes and Genomes (KEGG) through the KEGG Automatic Annotation Server (KASS) for functional annotation at the protein level [22]. We retrieved only 14% (738 out of 5,209) of the functional annotation for this set of genes, of which more than 59% (3,075 out of 5,209) comprises ORFs, encoding putative peptides shorter less than 100 aa in length. We explored the predominance of functions present in the 738 ORFs annotated at the KEEG Pathways level. Consequently, we found that in addition to proteins involved in more general functions, such as metabolic pathways (ko01100, 103 proteins) and the biosynthesis of secondary metabolites (ko01110, 30 proteins), proteins participating in more specific molecular tasks, such as the biosynthesis of antibiotics (ko01130, 22 proteins), the bacterial secretion system (ko03070, 20 proteins), ABC transporters (ko02010, 17 proteins), and two-component system (ko02020, 36 proteins), were frequently present as well. Among all of these proteins, we highlighted the presence of several members of the type II and IV secretion systems responsible for the secretion of bacterial toxins, proteins of the macrolide exporter system, and beta-lactamases and efflux pump proteins associated with beta-lactam resistance. Since such functional categories are found uniquely in different strains, this fact would support the idea that *P. aeruginosa* strains exhibit a wide variety of mechanisms to survive in several adverse environments being able to remain latently as reservoir of these genetic traits. Furthermore, this would have direct implication in emergence of multi-resistant and virulent strains since such genetic traits could all converge into single strains by horizontal transference mechanisms.

We further assessed the molecular functions of the portion of the *P. aeruginosa* accessory genome comprising genes between the 5th and 95th percentile of frequency (9 < accessory genome < 172) among all the genomes analysed. A total of 2,605 proteins were submitted again to the KASS server, retrieving functional annotation for 735 (28%) of them. We found a similar predominance of the above-stated pathways, but we expanded our analysis to include the biosynthesis of amino acids (ko01230, 37 proteins) and amino sugar and nucleotide sugar metabolism (ko00520, 13 proteins). Strikingly, we found additional proteins involved in vancomycin resistance as well as proteins of the type I and VI secretion systems associated with the export of toxins, proteases, lipases and other effector proteins. A general view of the molecular functions confined to different categories of the *P. aeruginosa* pangenome is shown in Figure 2. Comparison at the orthology level (Figure 2) indicated that a high level of functional specificity exists in all gene categories of the *P. aeruginosa* pangenome, whereby 79% of annotated genes in the core genome are not present in other categories. This percentage remains high at 47% in unique genes and 49% in the accessory genome. Previous studies have shown similar results in terms of functional categories of core and accessory genomes partially defined for *P. aeruginosa*, where core genome is enriched in central metabolism functions and major cellular functions such as replication, transcription, and translation as well as other associated biosynthetic pathways [19, 20]. At the KEGG functional module level, we disclosed some molecular pathways to be distinctive for every gene category in the pangenome. Table 2 summarizes those molecular pathways in which the *P. aeruginosa* core genome was found to contain a wide range of genes involved in either antibiotic biosynthesis and resistance. Therefore, functional characterization of the *P. aeruginosa* core genome would indicate that the infectivity and resistance are features intrinsically exhibited by any *P. aeruginosa* strain and that virulence and lethality would be confined to genetic traits encoded in the accessory genome.

**Figure 2.**
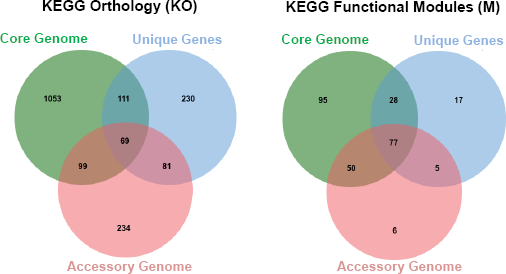
Functional annotation of the pangenome according to gene family categorization. Two Venn diagrams are presented, indicating the functional annotation at the orthology level (left diagram) and molecular pathway level (right diagram) for the three different categories established in concordance with gene frequency across strains. The redundancy of functions was predominantly at the pathway level and permitted to discern distinctive elements for each gene category. Those distinctive pathways are listed in Table 2.

**Table 2.** KEGG functional modules distinctive for the *P. aeruginosa* pangenome gene

| KEGG Module Number | Description | Genes |
| --- | --- | --- |
| Core Genome |  |  |
| M00064 | ADP-L-glycero-D-manno-heptose biosynthesis | 2 |
| M00493 | AlgZ-AlgR (alginate production) two-component regulatory system | 1 |
| M00235 | Arginine/ornithine transport system | 3 |
| M00531 | Assimilatory nitrate reduction | 1 |
| M00475 | BarA-UvrY (central carbon metabolism) two-component regulatory system | 2 |
| M00086 | Beta-Oxidation | 1 |
| M00123, M00573, M00577 | Biotin biosynthesis | 3 |
| M00364, M00366 | C10-C20 isoprenoid biosynthesis | 2 |
| M00170, M00171 | C4-dicarboxylic acid cycle | 2 |
| M00168 | CAM (Crassulacean acid metabolism) | 1 |
| M00722, M00727, M00728 | Cationic antimicrobial peptide (CAMP) resistance | 3 |
| M00256 | Cell division transport system | 1 |
| M00010 | Citrate cycle | 4 |
| M00120 | Coenzyme A biosynthesis | 3 |
| M00338 | Cysteine biosynthesis | 1 |
| M00154, M00155 | Cytochrome c oxidase | 3 |
| M00417 | Cytochrome o ubiquinol oxidase | 3 |
| M00552 | D-galactonate degradation, De Ley-Doudoroff pathway | 2 |
| M00596 | Dissimilatory sulfate reduction | 1 |
| M00542 | EHEC/EPEC pathogenicity signature | 2 |
| M00008 | Entner-Doudoroff pathway | 1 |
| M00445 | EnvZ-OmpR (osmotic stress response) two-component regulatory system | 3 |
| M00515 | FlrB-FlrC (polar flagellar synthesis) two-component regulatory system | 1 |
| M00729 | Fluoroquinolone resistance | 1 |
| M00344, M00345 | Formaldehyde assimilation | 3 |
| M00497 | GlnL-GlnG (nitrogen regulation) two-component regulatory system | 1 |
| M00605 | Glucose/mannose transport system | 2 |
| M00012 | Glyoxylate cycle | 1 |
| M00050 | Guanine ribonucleotide biosynthesis, IMP | 3 |
| M00259 | Heme transport system | 1 |
| M00045 | Histidine degradation | 4 |
| M00226 | Histidine transport system | 1 |
| M00620 | Incomplete reductive citrate cycle | 3 |
| M00131 | Inositol phosphate metabolism | 1 |
| M00190 | Iron(III) transport system | 4 |
| M00535 | Isoleucine biosynthesis | 3 |
| M00113 | Jasmonic acid biosynthesis | 1 |
| M00505 | KinB-AlgB (alginate production) two-component regulatory system | 2 |
| M00080 | Lipopolysaccharide biosynthesis | 1 |
| M00320 | Lipopolysaccharide export system | 2 |
| M00255 | Lipoprotein-releasing system | 1 |
| M00116 | Menaquinone biosynthesis | 1 |
| M00740 | Methylaspartate cycle | 2 |
| M00189 | Molybdate transport system | 3 |
| M00711 | Multidrug resistance, efflux pump MdtIJ | 1 |
| M00115 | NAD biosynthesis | 3 |
| M00144 | NADH:quinone oxidoreductase | 13 |
| M00471 | NarX-NarL (nitrate respiration) two-component regulatory system | 1 |
| M00622 | Nicotinate degradation | 1 |
| M00615 | Nitrate assimilation | 2 |
| M00438 | Nitrate/nitrite transport system | 1 |
| M00439 | Oligopeptide transport system | 1 |
| M00209 | Osmoprotectant transport system | 2 |
| M00004, M00007 | Pentose phosphate pathway | 6 |
| M00024 | Phenylalanine biosynthesis | 3 |
| M00434 | PhoR-PhoB (phosphate starvation response) | 1 |
| M00222 | Phosphate transport system | 4 |
| M00501 | PilS-PilR (type 4 fimbriae synthesis) two-component regulatory system | 1 |
| M00133 | Polyamine biosynthesis | 3 |
| M00015 | Proline biosynthesis | 3 |
| M00247, M00258 | Putative ABC transport system | 3 |
| M00193 | Putative spermidine/putrescine transport system | 7 |
| M00046 | Pyrimidine degradation | 1 |
| M00053 | Pyrimidine deoxyribonucleotide biosynthesis, CDP/CTP | 5 |
| M00052 | Pyrimidine ribonucleotide biosynthesis, UMP | 3 |
| M00377 | Reductive acetyl-CoA pathway (Wood-Ljungdahl pathway) | 1 |
| M00167 | Reductive pentose phosphate cycle | 5 |
| M00523 | RegB-RegA (redox response) two-component regulatory system | 2 |
| M00125 | Riboflavin biosynthesis, GTP | 2 |
| M00394 | RNA degradosome | 1 |
| M00308 | Semi-phosphorylative Entner-Doudoroff pathway | 2 |
| M00185 | Sulfate transport system | 2 |
| M00436 | Sulfonate transport system | 3 |
| M00435 | Taurine transport system | 1 |
| M00089 | Triacylglycerol biosynthesis | 2 |
| M00332 | Type III secretion system | 2 |
| M00025, M00040 | Tyrosine biosynthesis | 4 |
| M00117, M00128 | Ubiquinone biosynthesis | 7 |
| M00029 | Urea cycle | 3 |
| M00651 | Vancomycin resistance | 2 |
| M00241 | Vitamin B12 transport system | 1 |
| M00660 | Xanthomonas spp. pathogenicity signature | 2 |
| M00242 | Zinc transport system | 2 |
| Accessory Genome |  |  |
| M00502 | GlrK-GlrR (amino sugar metabolism) two-component regulatory system | 1 |
| M00533 | Homoprotocatechuate degradation | 2 |
| M00240 | Iron complex transport system | 3 |
| M00005 | PRPP biosynthesis | 1 |
| M00473 | UhpB-UhpA (hexose phosphates uptake) two-component regulatory system | 1 |
| M00644 | Vanadium resistance | 1 |
| Unique Genes |  |  |
| M00653 | AauS-AauR (acidic amino acids utilization) two-component regulatory system | 1 |
| M00500 | AtoS-AtoC (cPHB biosynthesis) two-component regulatory system | 1 |
| M00450 | BaeS-BaeR (envelope stress response) two-component regulatory system | 1 |
| M00104 | Bile acid biosynthesis | 1 |
| M00581 | Biotin transport system | 1 |
| M00569 | Catechol meta-cleavage | 4 |
| M00582 | Energy-coupling factor transport system | 1 |
| M00760 | Erythromycin resistance | 1 |
| M00524 | FixL-FixJ (nitrogen fixation) two-component regulatory system | 1 |
| M00713 | Fluoroquinolone resistance | 1 |
| M00059 | Glycosaminoglycan biosynthesis | 1 |
| M00499 | HydH-HydG (metal tolerance) two-component regulatory system | 1 |
| M00714, M00645 | Multidrug resistance | 2 |
| M00664 | Nodulation | 1 |
| M00549 | Nucleotide sugar biosynthesis | 1 |
| M00267 | PTS system, N-acetylglucosamine-specific II component | 1 |

### Molecular evolution in the Pseudomonas aeruginosa pangenome

In addition to uncovering the genes and functions that confer distinctive features to *P. aeruginosa* strains, we explored the genetic variability in every gene family retrieved from its pangenome. This approach could provide evidence of how the *P. aeruginosa* genome evolves to evade the immune response as well as depict the level of variability thought to be the major cause of the lack of success in designing a effective vaccine. For more than 10,000 gene families containing at least 2 members, we calculated the synonymous (dS) and non-synonymous (dN) rates, parameters indicative of the selection pressure on coding genes. The global distribution of dS and dN rates expressed as the omega value (*ω* = dN/dS) across the *P. aeruginosa* pangenome is presented in Figure 3A. Although the distribution of ω values fits well into a unimodal distribution, globally, it shows a shift-to-left distribution towards values lower than 1 with *ω* median = 0.1. These data suggest that the *P. aeruginosa* coding genome is under purifying selection as a whole, in which synonymous substitutions are predominantly higher than non-synonymous substitutions. The coding genes considered under positive selection must present *ω* > 1 (dN > dS); however, at the initial stage, we performed more restrictive filtering, thus considering those genes that exhibited at least a 2-fold greater non-synonymous substitution rate than the synonymous substitutions (*ω* ≥ 2). As a result, we retrieved a total of 230 genes (1.4% of pangenome) for which 71 functional annotations (31%) were recovered from the KASS server. We found a wide variability in terms of the molecular pathways for the genes under positive selection. Notably, among all genes under positive selection, we detected that some of them coded for proteins with remarkable functions, such as VirB2 and VirB9 (K03197 and K03204, respectively). Both proteins are components of the type IV secretion system and are localized at the outer membrane. In the case of VirB2 proteins, the T-pilus protein controls attachment to different receptors on the host cell surface to deliver toxin effector molecules [23]. Attempts to distinguish the specific role of these proteins through homologue searching in the Uniprot database have retrieved unclear results given that amino acid sequences of VirB2 and VirB9 from *P. aeruginosa* pangenome matched primarily with conjugal transfer proteins from *A. tumefaciens* (identity ~40% over 70% of the protein length), but also with toxin liberation protein F from *B. pertussis* (identity ~28% over 80% of the protein length). In any event, the VirB2 and VirB9 proteins must be exposed on the cell surface of pathogens making possible the *P. aeruginosa* be recognized by the host immune system and triggering a specific response against these potential antigens, thus promoting immune memory against this pathogen. The antigenicity of VirB2 and VirB9 proteins is further supported by their high rate of non-synonymous substitutions observed across different strains analysed, which would be result of the strong selection forces from the host immune system. Notwithstanding, we cannot discard these high rates of non-synonymous substitutions appear as response of phage predation. In this last scenario, the information retrieved in the present study regarding the set of genes under strong positive selection can be also useful to design bacteriophage-based therapies which have already been tested in *P. aeruginosa* [24]. Similarly, other outer membrane-bound proteins, such as the flippase MurJ (K03980) and the flagellin FlgF (K02391), which have been associated with virulence and pathogenicity [25, 26], exhibited a higher rate of non-synonymous substitutions than synonymous substitutions.

**Figure 3.**
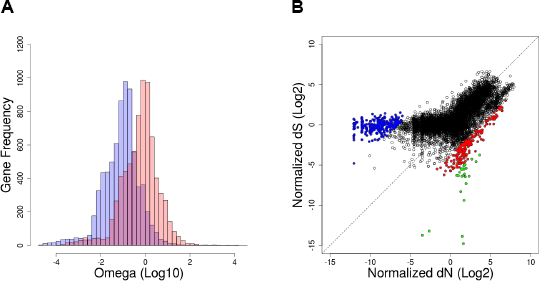
Molecular evolution of the *P. aeruginosa* pangenome. A – Histogram showing the distribution of omega (ω) values across the *P. aeruginosa* pangenome. The light blue histogram shows the original distribution with the tendency towards values indicating purifying selection (shift to left from neutrality). The superposed light red histogram indicates the Z-scores for the selection of genes with ω significantly different than 1. Those with significant ω < 1 were considered to be under strong purifying selection for functional analysis, and those with significant ω > 1 were selected to be under strong positive selection for the MLST approach. B – Scatter plot to represent the distribution of normalized dN and dS rates for all gene families detected in the *P. aeruginosa* pangenome. Gene families under strong purifying selection are highlighted in blue, whereas gene families under positive selection (ω > 2) are highlighted in red. The set of gene family candidates for MLST under strong positive selection are highlighted in green. The diagonal dashed line indicates the boundary for neutrality.

Strong selection forces from the immune response or environmental pressure were also detected in a set of *P. aeruginosa* genes tightly linked with virulence in other human pathogens. Therefore, we observed positive selection in the following genes: the PsrtC (K08303) homologue, a protease involved in mucus degradation during *H. pylori* infection (pathway ko05120); the MprF and ParR homologues (KO14205 and K18073, respectively), proteins involved in the cationic antimicrobial peptide (CAMP) resistance in Gram-positive and Gram-negative bacteria (ko1503), respectively; the PstS homologue (K02040), an integral membrane ABC phosphate transporter that modulates the TLR4 response during *M. tuberculosis* infection (ko5152); the *T. brucei* ICP homologue (14475), a protein involved in immunosupression by modulating the degradation of IgGs (ko5143); and the RNA polymerase sigma-54 factor (K03092), which is associated with the *V. cholera* pathogenic cycle to control the expression of motor components of flagella (ko5111).

Given the low level of functional annotation for genes under positive selection, we performed an additional quantitative assessment to determine protein domain enrichment in the group of proteins under positive selection using the Simple Modular Architecture Research Tool (SMART) and the Protein Family database (Pfam) nomenclature systems. Once the inventory of SMART and Pfam domains contained in the entire *P. aeruginosa* pangenome was assessed, we performed a Fisher’s exact test for 2 x 2 contingency tables to verify the significant over-representation of Pfam/SMART domains in the proteins under positive selection with respect to the pangenome. We observed the presence and prevalence of 4,090 different protein domains from both the SMART and Pfam classification in the *P. aeruginosa* pangenome. Forty-four of these 4,090 domains were found to be over-represented in the proteins exhibiting positive selection (Table 3). Among them, we observed a high frequency of membrane-bound proteins acting as transporters or receptors. Some of the functions over-represented in Table 3 agree with some stated from previous analyses in which membrane proteins (transporters and/or receptors) as well as the Sigma-54 factor seem to be under positive selection in *P. aeruginosa*. Interestingly, we observed the presence of proteins related with either 16S RNA and ribosomal protein methylation (Table 3). We detected such patterns of molecular evolution in this class of proteins previously, but in different human pathogens [27]. Although we cannot shed light on the meaning of this type of evolution in these proteins given their function, we hypothesized that they might influence the translation process to modulate the expression of a certain set of proteins directly or indirectly involved in pathogenesis. Recent studies on rRNA methylation indicate that they play a meaningful role in decoding function [28–30]. Indeed, some of them have been directly involved with virulence [31].

**Table 3.** Domain enrichment in proteins under positive selection

| SMART/Pfam Domain | Description | Fisher's test |
| --- | --- | --- |
| Chromate_transp | Probably act as chromate transporters in bacteria | 0.0000 |
| Sulfatase | Present in esterases hydrolysing steroids, carbohydrates and proteins | 0.0020 |
| PepSY_TM | Conserved transmembrane helix found in bacterial protein families | 0.0041 |
| PrmA | Present in the Ribosomal protein L11 methyltransferase | 0.0123 |
| Cons_hypoth95 | Present in 16S RNA methyltransferase D | 0.0166 |
| MTS | Present in the 16S RNA methyltransferase C | 0.0182 |
| DUF1329 | Putative outer membrane lipoprotein | 0.0215 |
| DUF4102 | Putative phage integrase | 0.0235 |
| CHASE | Extracellular domain of bacterial transmembrane receptors | 0.0284 |
| G3P_acyltransf | Enzymes converting glycerol-3-phosphate into lysophosphatidic acid | 0.0284 |
| AceK | Bacterial isocitrate dehydrogenase kinase/phosphatase protein | 0.0284 |
| Choline_sulf_C | C-terminus of enzyme producing choline from choline-O-sulfate | 0.0284 |
| DUF2165 | Unknown function | 0.0284 |
| DUF2909 | Unknown function | 0.0284 |
| DUF3079 | Unknown function | 0.0284 |
| DUF444 | Unknown function | 0.0284 |
| DUF533 | Unknown function; integral membrane protein | 0.0284 |
| DUF791 | Unknown function | 0.0284 |
| DUF972 | Unknown function | 0.0284 |
| Glu_cys_ligase | Enzyme carrying out the first step of glutathione biosynthesis | 0.0284 |
| Herpes_UL6 | Present in proteins similar to herpes simplex UL6 virion protein | 0.0284 |
| His_kinase | Membrane sensor, a two-component regulatory system | 0.0284 |
| Inhibitor_I42 | Protease inhibitor | 0.0284 |
| PPDK_N | Present in enzymes catalysing the conversion of pyrophosphate to PEP | 0.0284 |
| Sigma54_AID | Activating interacting domain of the Sigma-54 factor | 0.0284 |
| Sigma54_CBD | Core binding domains of the Sigma-54 factor | 0.0284 |
| Sigma54_DBD | DNA binding domain of the Sigma-54 factor | 0.0284 |
| PAS, PAS 4/9 | Present in signalling proteins working as signal sensors | 0.0330 |
| MFS | Major Facilitator Superfamily of small molecule transporters | 0.0359 |
| Autoind_synth | Autoinducer synthase involved in quorum-sensing response | 0.0423 |
| AzIC | Putative protein involved in branched-chain amino acid transport | 0.0423 |
| Chitin_bind | Present in carbohydrate-active enzymes (glycoside hydrolases) | 0.0423 |
| DUF3299 | Unknown function | 0.0423 |
| PTS_EIIC / IIB | Phosphoenolpyruvate-dependent phosphotransferase system | 0.0423 |
| TctC | Member of the tripartite tricarboxylate receptors | 0.0423 |
| UPF0004 | Domain found in tRNA methyltransferases | 0.0423 |

When we attempted a similar analysis in a counterpart set of proteins under purifying or negative selection (*ω* < 1), the biased distribution of omega values across the *P. aeruginosa* pangenome (Figure 3A) made it difficult to set up a suitable threshold to recover proteins under this type of selection. Therefore, we obtained Z-scores of both the dN and dS rates (Figure 3A, light red histogram), thus reaching a normal distribution around *ω* = 1 (neutrality). Using this normalized distribution of *ω* values, we could determine those genes with evolution significantly different (p ≤ 0.05) from neutrality (*ω* = 1) towards a strong negative selection (lowest *ω* values). As a result, we found a group of 268 proteins/genes under negative selection, the dN and dS rates of which are plotted in Figure 3B (see the blue points distribution). The quantitative assessment to determine protein domain enrichment indicated that more than 130 SMART and/or Pfam domains were over-represented in this set of proteins, and as expected, most of them were related to the central functions of cell maintenance, such as translation (ribosome proteins, tRNA biogenesis, amino acid starvation response), carbohydrate metabolism, amino acid biosynthesis and transport, and respiration.

*New high variability markers for multi-locus sequence typing of P. aeruginosa strains* Characterization of the *P. aeruginosa* pangenome offers not only critical information about the molecular functions and prevalence of certain genes across multiple strains analysed but also information about the level of genetic variability at the strain level. A molecular evolution approach retrieved a large set of genes/proteins under positive selection in *P. aeruginosa*. At the same time, such genes could be used for genotyping aims to associate certain genetic variants with pathogenicity and virulence traits. As a consequence, we selected and tested some *P. aeruginosa* genes in a MLST strategy to discern phylogenetic relationships among a large number of PATRIC reference strains analysed and six *P. aeruginosa* aminoglycoside and carbapenem-resistant strains isolated from patients who acquired healthcare-associated infections in a clinic located outside the metropolitan area of Medellin, Antioquia, Colombia.

We narrowed down the list of MLST candidates by selecting the genes that had the following characteristics: i) present in at least 95% of the strains explored at the sequence level (frequency ≥ 172); ii) exhibiting omega values significantly higher than 1 (Figure 3B, p ≤ 0.05, ω > 15); and iii) short enough to facilitate Sanger sequencing in a few reactions. Of the 27 genes/proteins showing significant positive selection, we finally selected four genes, the features of which are depicted in Table 4. After amplification and Sanger sequencing of selected genes in our six *P. aeruginosa* isolates, we combined that genetic information with that completely available for 170 *P. aeruginosa* strains, thus building a multiple sequence alignment almost 3,000 bp in length for a total of 176 different strains. Using maximum likelihood approaches, we reconstructed the phylogenetic relationships among all strains and retrieved the phylogenetic tree showed in Figure 4. Our six local isolates were positioned in three different clades, where isolate 49 was closely related to the highly virulent *P. aeruginosa* PA14 strain, representing the most common clonal group worldwide [32]. By contrast, isolate 77 was related to several strains, including the multi-drug-resistant *P. aeruginosa* NCGM2.S1 [33] and the cytotoxic corneal isolate *P. aeruginosa* 6077 [34]. Finally, the 30-1, 42-1, 45, and 04 isolates presented a close relationship and were related to the multi-drug resistant *P. aeruginosa* VRFPA02 isolate from India [35].

**Figure 4.**
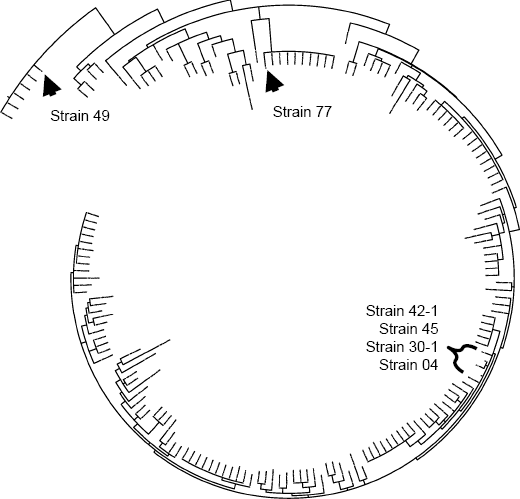
Circular phylogenetic tree showing the genetic relationships among 170 reference PATRIC strains and our six *P. aeruginosa* isolates. The phylogenetic tree was built from the best evolutionary model explaining evolution at the concatenated gene families 3333, 3675, 4766, and 5348 after a sequential likelihood ratio test [54]. A total of 176 *P. aeruginosa* strains are located in the tree, and the localization of our clinical isolate is indicated. A close view of this tree permitted us to infer relationships among our clinical isolates with virulent and multi-drug resistant strains.

**Table 4.** Potential genetic markers for MLST in *P. aeruginosa* strains

| Gene<br>Family<sup>a</sup> | Function<sup>b</sup> | Omega<br>($\omega$) | Length<br>(bp) | Strain<br>Frequency<sup>c</sup> |
| --- | --- | --- | --- | --- |
| 3333 | Chitin binding protein | 108 | 1,170 | 98.9% (179) |
| 3675 | Flagellar basal-body rod protein FlgF | 5,884 | 750 | 99.5% (180) |
| 4766 | Predicted branched-chain amino acid permease AzIC | 86 | 763 | 96.7% (175) |
| 5348 | Unknown function | 32 | 573 | 99.5% (180) |

a Nomenclature according to pangenome gene inventory.
b Function inferred from KEGG, SMART, and/or BLAST-based search.
c Number of strains carrying respective genes are denoted in parenthesis.

Based on the best evolutionary model fitted to the nucleotide substitution pattern observed for these markers (TrN+I+G), a proportion of invariable sites of 0.9080 was obtained, thus indicating that more than 250 polymorphic sites are present in our MLST approach. Moreover, gamma distribution parameters (0.5060) is indicative of few hot-spots with high substitution rates [36]. In this fashion, we provided support to use the highly variable genetic markers reported here for MLST to produce an initial, fast, and cost-effective genotyping for *P. aeruginosa* strains of clinical interest. To compare if the evolutionary history of *P. aeruginosa* strains is equally represented by of our proposed MLST markers in comparison with that inferred by using common MLST markers [37, 38], we reconstructed a phylogeny using similar approaches and DNA sequences corresponding to seven housekeeping genes: *acsA, aroE, guaA, mutL, nuoD, ppsA*, and *trpE*. The resulting tree showed not deep topology differences when compared to that created from our proposed MLST approach (data not shown). This indicate that the new molecular markers proposed in this study for genotyping aims could be used to infer the evolutionary history of *P. aeruginosa* strains.

## Conclusions

High-throughput sequencing technology has permitted the analysis of the genetic identity of a vast number of microorganisms, an applied science especially relevant to studying human pathogens and their virulence and pathogenicity traits in depth. Here, we have performed a reverse vaccinology approach using a large amount of genetic information available in the PATRIC database to determine the genetic elements of *Pseudomonas aeruginosa* to be probably targeted in future clinical studies aiming new vaccine designs. We have extensively described the *P. aeruginosa* pangenome in terms of the effective number of non-redundant genes present in this bacterial species by analysing more than 180 different strain genomes. We outlined the genomic variability of this human pathogen, demonstrating that approximately 60% of the *P. aeruginosa* genome is variable across strains, with the remaining genome encoding genes that are involved in central functions, such as virulence, resistance, toxicity and pathogenicity.

We have identified major genetic pieces of the core and accessory genome in *P. aeruginosa*. Approximately 15% (2,503/16,820 genes) of the pangenome was found to constitute the core genome and was present in 100% of the strains studied, accomplishing general molecular functions for cell maintenance such as replication, translation, transcription, central metabolism, electron transport chain, amino acid biosynthesis and transport, nucleotide biosynthesis, cell wall synthesis and maintenance, and cell division. Conversely, the accessory genome exhibited a comprehensive variety of functions, ranging from a wide spectrum of antibiotic resistances to a specialized secretion system delivering toxins and effector proteins potentially harmful for host cells. However, pathogenicity traits were also observed in the distinctive KEGG pathways revealed for the core genome.

Although this is not the first report to describe the pangenome for a single bacterial species [14, 21, 39, 40], and other very recent studies have attempted to determine the *P. aeruginosa* pangenome [18–20], this report is the first to describe a closed *P. aeruginosa* pangenome at very large scale, including almost 200 bacterial genomes from this human pathogen and performing a pangenome-scale molecular evolutionary analysis. Our study fits well with previous and general genomic characterizations of this human pathogen [18–20, 41], and it definitely expands our knowledge about the evolutionary mechanisms of *P. aeruginosa* pathogenesis. This study aimed to reveal the evolutionary processes occurring at the pangenome level in *P. aeruginosa* that could explain the failure to design and develop of a successful vaccine against this human pathogen as well as provide an understanding of the molecular mechanisms that drive the evasion of the host immune system. We observed that the *P. aeruginosa* genome is globally under purifying selection, given the distribution of omega values (*ω* = dN/dS, median ~0.1) discerned for every gene family present in its pangenome. This result was further supported by the finding that there are 10-fold more genes under strong purifying selection than strong positive selection (significantly different to neutrality, p ≤ 0.05). Although we found that the *P. aeruginosa* pangenome evolves to purifying selection as a whole, we distinguished some genes and functions predominantly present in the reduced set of genes under positive selection. As a consequence, a considerable number of proteins located at the outer membrane, such as those associated with receptor and transporter functions, were identified to have an increased rate of non-synonymous substitutions. These data corroborated our results based on KEGG functional analysis, which described an ample group of surface-exposed proteins under strong selection forces from the immune response or environmental pressure.

For the first time, pangenome-scale evolutionary information is presented to support the design of new *P. aeruginosa* vaccines. In this fashion, failures when using protein conjugates from pili, flagella, or secretions systems [5, 7, 9, 11] are partially explained by the data presented here, which indicates the presence of a high genetic variation in this class of proteins in terms of non-synonymous substitutions, a fact that has been described previously but at very lower scale [42, 43].

Finally, we further explored the genetic information derived from our molecular evolution analyses and proposed a set of four new polymorphic genetic markers for MLST. We demonstrated that these markers contain an adequate proportion of hotspots for variation, exhibiting high nucleotide substitution rates. Using these four loci, we discerned the genetic identity of 6 local isolates of *P. aeruginosa* and related them with the resistance and virulence traits carried in reference strains.

## Methods

### Pangenome-wide analysis

Genome information from *P. aeruginosa* strains was downloaded via the ftp server from the PATRIC database [13]. A set of 181 available genomes (*ffn* files) was retrieved from the PATRIC database, April 2014 release. Estimation of the *Pseudomonas aeruginosa* pangenome size was assessed in a similar manner to that previously reported as genome-and gene-oriented methods using iterative and combinatorial approaches [14, 15, 17]. Briefly, a BLAST-based iterative method was used to extract the full set of non-redundant genes representing the *P. aeruginosa* pangenome. A single iteration consisted in a random selection of a strain as pangenome primer, then the remaining set of strain were randomly incorporated to the pangenome. The above process was calculated over 200 iterations with random permutation of the strain order in every iterative step. A rarefaction curve was plotted with all data generated and consisted in 200 different measures throughout a sequential addition of 181 different strains. Pangenome metrics was also obtained in iterative manner with data fitting to the power law as previously stated [15]. Power regression was calculated individually for each iteration of pangenome reconstruction (n=200) and plotted in R v.3.1.2 with the *“igraph”* package (https://cran.r-project.org). Alpha parameter from the n = KN^-α^ power regression, indicating whether pangenome is open or closed, was calculated individually with least squares fit of the power law to the number of new genes discovered at core genome according to tettelin and coworkers [15]. Therefore, the global alpha value for the *P. aeruginosa* pangenome was determined as the mean of all 200 different alpha values generated ± sd with the confidence interval at 0.95 level. Finally, the set of non-redundant genes obtained was used to explore their occurrence pattern in the 181 *P. aeruginosa* genomes through BLASTN-based comparisons [44, 45].

### Molecular evolution analysis

The full set of ORFs constituting the *P. aeruginosa* pangenome was used to search homologues in all genomes analysed, and multiple sequence alignments were built using refined and iterative methods [46, 47]. The synonymous and non-synonymous substitution rates were calculated in a pairwise manner using approximate methods [48] and by correcting for multiple substitutions [49]. Omega values (*ω*) were computed as the averaged ratio of dN/dS rates from multiple comparisons, and genes under strong positive selection were selected when *ω* ≥ 2. The Z-score of ω values was computed to depict functions of genes under strong purifying selection and potential MLST genetic markers under strong positive selection (p ≤ 0.05). Large-scale analyses of pairwise comparisons, statistical analysis, and graphics were performed using R v3.1.2 (https://cran.r-project.org).

### Functional genomics analysis

Functional annotation of genes was performed using the KEGG Automatic Annotation Server for KEGG Orthology [22]. KEGG functional modules and ontologies were explored in the KEGG BRITE database [50]. Functional domains present in genes of interest were assigned using Perl scripting for batch annotation (http://smart.embl-heidelberg.de/help/SMART_batch.pl) against the Simple Modular Architecture Research Tool (SMART) together with Pfam classification [51, 52]. Fisher’s exact test with a false discovery rate (FDR) for 2 x 2 contingency tables to measure enrichment of Pfam/SMART domains was performed using R v3.1.2 (https://cran.r-project.org). Venn diagrams were drawn using the *jvenn* server [53].

### Multi-locus sequence typing

The six *P. aeruginosa* strains (labelled as 04, 30-1, 42-1, 45, 49, and 77) were isolated from patients who acquired healthcare-associated infections at a clinic located outside the metropolitan area of Medellin, Antioquia, Colombia. This study was approved by the ethics committee of the Fundación Clínica del Norte Hospital (Bello, Antioquia, Colombia). The six isolates, previously characterized for multi-drug resistance, were kindly donated to the scientist of the Bacteria & Cancer Researching Group of the Faculty of Medicine, University of Antioquia, Colombia. The genomic DNA from *P. aeruginosa* multi-drug resistant strains was extracted using a GeneJER™GenomicDNA Purification Kit (Thermo Scientific, Waltham, MA, USA). The reference sequences of *P. aeruginosa* PA01 for the four markers selected to perform MLST were downloaded from a public database [GenBank: AE004091.2: region 930623 to 931822 (family 3333), region 1167488 to 1168237 (family 3675), region 2230183 to 2229425 (familyc 4766), region 2935851 to 2936423 (family 5348)]. Primers were designed to amplify the complete sequence of each gene, and the Polymerase Chain Reaction (PCR) proceeded with 28 cycles of amplification using Phusion^®^ High-Fidelity DNA Polymerase (Thermo Scientific, Waltham, MA, USA) and 50 ng of genomic DNA. PCR products were isolated using a GeneJet PCR Purification Kit (Life technologies, Carlsbad, CA, USA), and both strands were sequenced by the Sanger automatic method in an ABI 3730 xl instrument (Stab Vida Inc., Caparica, Portugal). Base calling and genetic variants were manually explored using the delivered *ab1* files and FinchTV viewer (Geospiza Inc. Perkin Elmer, Waltham, MA, USA). Assembled sequences from both strands were obtained and concatenated to respective reference sequences obtained from the PATRIC genomes analysed. Sequences belonging to the respective gene family were aligned using iterative methods [46, 47], and alignments were concatenated to perform phylogenetic analysis. The sequential likelihood ratio test was carried out to detect the evolutionary model that better explained genetic variation in all genes integrated in the MLST approach. For that reason, we used the jModelTest tool [54], and model selection was completed by calculating the corrected Akaike Information Criterion (cAIC). The MLST tree was constructed using the Interactive Tree Of Life (iTOL) tool [51, 55] and the phylogeny obtained using the TrN+I+G model. For comparisons aims, we compiled genetic information from seven MLST markers commonly used in *P. aeruginosa* genotyping being the housekeeping genes: *acsA, aroE, guaA, mutL, nuoD, ppsA*, and *trpE* [37, 38]. Aligned sequences were concatenated and phylogenetically analyzed with the jModelTest tool as well. Tree topology generated from this conventional MLST markers was compared with that obtained using the new MLST markers proposed in this study.

## Availability of supporting data

The features of the Pseudomonas aeruginosa strains used in this study are included in the Additional File 1. All the DNA sequences derived from PCR amplification and Sanger sequencing of the four MLST studied here for the *P. aeruginosa* clinical isolates were submitted to the GenBank through BankIt server [GenBank: KU214214 to KU214237].

## Competing interests

The authors declare that they have no competing interests.

## Authors’ contributions

ABP designed and directed this study. JMR and ABP performed the pangenome, molecular evolution, and phylogenetic analyses. ERP and AMR obtained the *P. aeruginosa* clinical isolates. JMR, AMR, and MC performed PCR techniques. JMR and SCB curated the sequences from Sanger automatic sequencing. JMR, SCB, and ABP prepared the manuscript. All authors read and approved the final version of this manuscript.

## Acknowledgements

The authors give thanks to the Colombian Agency for Science, Technology, and Innovation (Colciencias) and the National Fund for Science, Technology, and Innovation “Francisco José de Caldas” for grant 5817-5693-4856 to ABP and grant 1115-5693-3375 to ERP. The authors also thank the “Clinica Antioquia” microbiology laboratory staff, who donated the clinical isolates for the MLST studies. The JMR M.Sc. fellowship was supported by the Colombian Agency for Science, Technology, and Innovation (Colciencias) with funds of the 5817-5693-4856 grant.

Catalogue of the KEGG functional modules (M) distinctively found in three gene categories of the *P. aeruginosa* pangenome: core, accessory, and unique genes. The number of modules correlated with those numbers presented in Figure 2 (Venn diagram on the right).

The SMART and Pfam domains are presented in a non-redundant manner. Function description was recovered from annotations in SMART or Pfam databases. Fisher’s test values correspond to p-values (p ≤ 0.05), supporting the over-representation of the corresponding domain in the set of proteins under positive selection.

